# Prevalence and global distribution of *bla_KPC-2_* and *bla_NDM-1_* genes in *Klebsiella Pneumoniae*

**DOI:** 10.1101/2020.02.18.955211

**Authors:** Xiufeng Zhang, Fangping Li, Shiyun Cui, Lisha Mao, Xiaohua Li, Awan Furqan, Weibiao Lv, Zhenling Zeng

**Affiliations:** Guangdong Provincial Key Laboratory of Veterinary Drugs Development and Safety Evaluation, South China Agricultural University, Guangzhou 510642, People’s Republic of China; National Laboratory of Safety Evaluation (Environmental Assessment) of Veterinary Drugs, South China Agricultural University, Guangzhou 510642, People’s Republic of China; National Risk Assessment Laboratory for Antimicrobial Resistance of Animal Original Bacteria, South China Agricultural University, Guangzhou 510642, People’s Republic of China; Guangdong Provincial Key Laboratory of Plant Molecular Breeding, State Key Laboratory for Conservation and Utilization of Subtropical Agro-Bioresources, South China Agricultural University, Guangzhou 510642, People’s Republic of China; Department of Clinical Laboratory, Cancer Hospital of Guangxi Medical University, Guangxi Medical University, Nanning, 530021, People’s Republic of China; Department of Clinical Laboratory, Shunde Hospital, Southern Medical University (The First People’s Hospital of Shunde), Foshan 528000, People’s Republic of China

**Keywords:** *Klebsiella pneumonia*, *bla_KPC-2_*, *bla_NDM-1_*, Bioinformatics

## Abstract

Carbapene-resistant *Klebsiella pneumoniae* infections have caused a major concern and posed a global health threat to public. *bla_KPC-2_* and *bla_NDM-1_* genes are the most widely reported of carbapenem resistance genes in *K. pneumoniae*. In this study, we investigated phylogenetic relationships of carbapene-resistant *K. pneumoniae* from a tertiary hospital between 2013 and 2018 in China and analyzed the global epidemiology and distribution of *bla_KPC-2_* and *bla_NDM-1_* gene in *K. pneumoniae* based on 1579 NGS genomes. We found that 19 carbapene-resistant *K. pneumoniae* isolated were divided into five lineages and all had high genotypic and phenotypic resistance. Two lineages (mostly ST11 and ST25) were the major type detected carrying *bla_KPC-2_* and *bla_NDM-1_* gene, respectively. Among global genomes data, 147 known ST types have been identified and ST11 and ST258 were the globally prevalent clones. Genetic environment analysis showed that the *ISKpn27-bla_KPC-2_-ISKpn6* and *bla_NDM-1_-ble-trpf-nagA* may be the core structure in the horizontal transfer of *bla_KPC-2_* and *bla_NDM-1_*, respectively. In addition, DNA transferase (*hin*) may be involved in the horizontal transfer or the expression of *bla_NDM-1_*. This study sheds some light on the genetic environment of *bla_KPC-2_* and *bla_NDM-1_* and should foster further studies about the mechanism of carbapene-resistant *K. pneumoniae* dissemination.

## Introduction

*Klebsiella pneumonia*, as a member of ESKAPE pathogens, is the most common factor of nosocomial infections. Infections caused by *K. pneumonia* are associated with a high risk of mortality and increased economic costs^1^. In recent years, *K. pneumonia* has been considered as a growing global threat due to its high level of resistance, particularly to those last resort antibiotics, such as carbapenems. In 2017, carbapenem-resistant *K. pneumonia* (CRKP) has been listed in the critical priority tier and the highest priority in new antibiotic development by WHO^2^.

Carbapenem resistance in *K. pneumoniae* involves multiple mechanisms, including the production of carbapenemases, alterations in outer membrane permeability and the upregulation of efflux systems. The latter two mechanisms are often combined with high levels of other types of β-lactamases (e.g., AmpC, ESBLs)^3^. Carbapenemases are carbapenem hydrolyzing β-lactamases and the most popular carbapenemases are ambler molecular class A (KPC), class B (VIM, IMP, NDM) and class D (OXA-48-like) types among *K. pneumoniae^4^*.

The KPC-type β-lactamases have been almost exclusively reported in *K. pneumonia*. Even though there are more than 20 different KPC variants reported, KPC-2 and −3 remain the most commonly identified variants^5^. The class B β-lactamases identified in *K. pneumonia* have been identified in various *Enterobacterial* species and worldwide^6^. The majority of NDM-1-producing *K. pneumonia* strains also carry a diversity of other resistance mechanisms but remain mostly susceptible to agents such colistin, fosfomycin, and tigecycline. The class D OXA-48-producing *K. pneumoniae* is endemic in Turkey and certain North African and European countries, and shows a wide range of susceptibility profiles^7^. These different carbapenemase genes circulating within *K. pneumoniae* are often carried by mobile structures, including insertion sequences (IS), plasmids and transposons, and therefore cause a significant public health concern^8^.

Nosocomial outbreaks caused by CRKP have been rapidly emerging in many countries and they are more often found in critically ill patients dwelling in intensive care units (ICU)^9^. Extensive genotypic characterization has shown that clusters of highly similar *K. pneumoniae* occur, representative of distinct clonal lineages and there are several high-risk clones or ST types reported for the global distribution of CRKP^10^. To date, the pandemic of KPC-producing *K. pneumoniae* is driven primarily by the spread of clonal complex 258 (CC258) isolates^11^. CC258 consists of the predominant clone ST258 and its single-locus variants (ST11, ST340 and ST512). Previous study indicated that *K. pneumoniae* ST11 is an emerging high-risk clone that is associated with *bla_KPC_*, ST14, ST25 and ST340 with *bla_NDM-1_* have been reported widely^12^.

Recently, carbapenem-resistant hypervirulent *K pneumoniae* strains have been reported and caused a substantial threat to human health because they are simultaneously hypervirulent, multidrug resistant, and highly transmissible^13,14^. Hence, it is of great significance to carry out molecular mechanism research and epidemiological investigation on carbapenem resistance genes among *K. pneumonia* in the world. With the development of sequence technology, there were *K. pneumonia* genomes available in the GenBank databases from NCBI and make it possible to study the evolution and epidemiology of carbapenem resistance in this *K. pneumonia* pathogen^15,16^.

In this study, we conducted a six-year-long longitudinal study of a tertiary care hospital in China, to study the time dynamics of carbapenem-resistant *K. pneumoniae* and presence of carbapenem resistance genes *bla_kPC-2_* and *bla_NDM-1_* on healthcare surfaces using next generation sequencing (NGS) and bioinformatic analyses. In addition, we combine the NGS *K. pneumoniae* data from the GenBank databases to demonstrate the *K. pneumoniae* distribution characteristics in the world, and to analyze the global distribution and gene structure characteristics of genes *bla_kPC-2_* and *bla_NDM-1_*. The study aims to provide theoretical basis for the rational use of clinical antibiotics and the reduction of the outbreak of nosocomial infection.

## Materials and methods

### Strains collection and Antimicrobial susceptibility

CRKP isolates were collected from Shunde hospital in China between 2013 and 2018. For all isolates, the minimum inhibitory concentrations (MICs) of antibiotics were determined using a BD PhoenixTM 100 Automated Identification and Susceptibility Testing system (BD, USA) according to CLSI and EUCAST guideline^17,18^.

### Illumina Whole Genome Sequencing

Bacterial genomic DNA of CRKP strains were extracted using a HiPure Bacterial DNA Kit (MAGEN). The libraries were created using the VAHTSTM Universal DNA Library Prep kit for Illumina. Whole genome sequencing was carried out on an Illumina Hiseq 2500 system to obtain 2×150 bp reads. Processed reads were de novo assembled into contigs with CLC Genomics Workbench 10.1 (CLC Bio, Aarhus, Denmark) and the genomes were annotated by NCBI Prokaryotic Annotation Pipeline (PGAP).

### Global data gathering

Data from the national center of biological information (NCBI) and its open database (https://www.ncbi.nlm.nih.gov/assembly), we used conventional python scripts to screen samples with complete information for further analysis. A total of 1579 samples from 44 countries, including 247 fully assembled level samples and 1332 scaffold level samples, were gathered to subsequent analysis.

### Taxonomic assignment and ARG, VG, IS identification

FastANI v1.1 (https://github.com/ParBLiSS/FastANI) with the core algorithm of BLAST-based ANI solver was used to identify the species of all isolates^19^. Genome ASM24018v2 (https://www.ncbi.nlm.nih.gov/search/all/?term=%20ASM24018v2%20) was used as *K. pneumoniae* reference genome for species identification. Species were determined if the genome in question had >95% ANIb and all the isolates from hospital have >98% ANIb compared with reference genome^20^. Software MLST v2.16.1 was used to perform multi-site sequence typing (MLST) on sample genome^21^. ResFinder BLAST identification program (https://cge.cbs.dtu.dk/services/ResFinder/) and Isfinder were used to identify acquired antibiotic resistance genes (ARGs) and mobile elements, respectively^22^. VFDB (http://www.mgc.ac.cn/VFs/) virulence database (setB) was conducted for bacterial virulence genes analysis.

### Clustering and Pan-genome analysis

Prokka v1.14 was used to produce gff file for the contig of *K. pneumoniae* genome we collected^23^. After that, core genome alignment was constructed with Roary v3.8.0 and PRANK v1.0^24^.

Core_genome_alignment.aln, the output file of Roary pipeline was conveyed to fastGEAR to identify Lineages by hierBAPS^25^.

### *Bla_KPC-2_* and *bla_NDM-1_* loci annotation and comparison

Regular Python scripts and Easyfigv2.2.3 (http://easyfig.sourceforge.net/) used in the gene sequence extracted for environment analysis.

Seventy-six *K. pneumoniae* genomes and thirty *K. pneumoniae* genomes with complete chromosomal genomic data were selected for *bla_KPC-2_* and *bla_NDM-1_* gene environment analysis, respectively. Through python script, we extracted the upstream and downstream 2~6 kb sequences of the target gene for blast alignment and annotation.

### Visualization of drug resistance and global distribution

ArcGIS 10.3, R package ggplot2 (http://had.co.nz/ggplot2), package pheatmap (https://stat.ethz.ch/pipermail/r-help/2012-November/330785.html) and conventional Python scripts were used for visualizing analysis.

### Nucleotide Accession Number

These assemblies sequence data of CRKP isolated from hospital were deposited in the GenBank database under BioProject accession PRJNA564463.

## Results

### CRKP isolates had high genotypic and phenotypic resistance

A total of 19 CRKP strains were isolated from Shunde hospital. All isolates resulted multi-resistant phenotypes and carried 29 unique ARGs for 8 different classes of antibiotics. All strains were sensitive to polymyxins and tigecycline (figure.1(b)). 94.7% of these ARGs were ß-lactamases and *bla_SHV_, bla_NDM-1_, bla_TEM-1B_, bla_OXA-1_, bla_KPC-2_* were the most prevalent for carbapenemase-encoding genes. In addition, NDM-producing strains contained more antibiotic resistance genes than KPC-producing isolates.

**Fig. 1.**
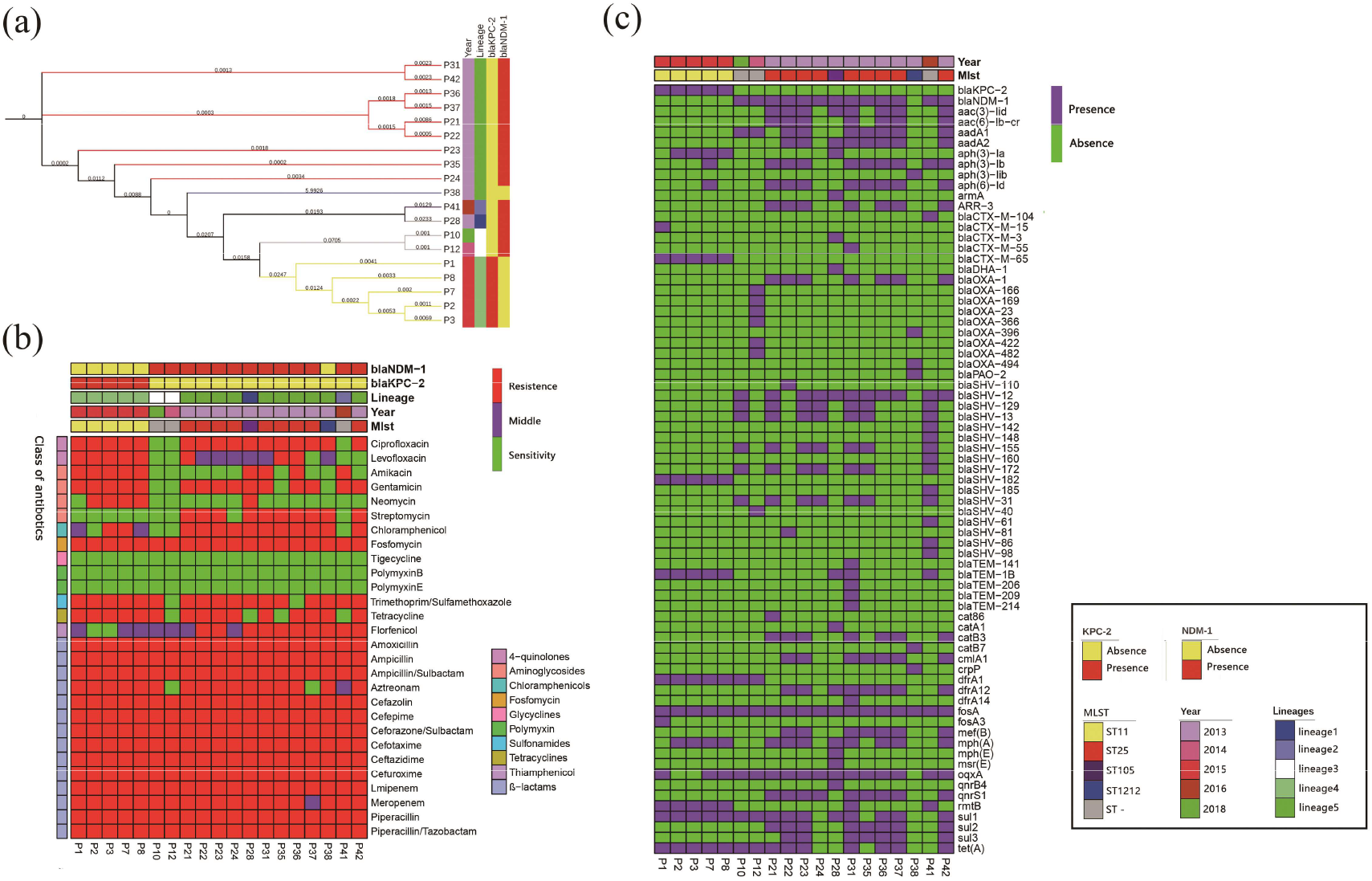
Results of phylogenetic trees; bayesian classification, MIC test and ARGs identification of 19 samples from ICU rooms. Sampling time, lineages of hierBAPS, presence/absence of *bla_KPC-2_* and *bla_NDM-1_* are annotated as colored bars next to the isolate number. (a)Phylogenetic trees of high abundance species from core genome alignments. Maximum likelihood phylogenetic trees from core genome alignments; Tree branches are colored by MLST types. (b) Results of MIC test; class of antibiotics and resistance strength are annotated as colored bars next to the main heatmap. (c) Results of ARGs identification; presence/absence of ARGs are annotated as colored bars next to the main heatmap.

### Two lineages dominated CRKP populations

MLST typing showed that CRKP isolates were divided into four ST types and three untyped ST. Most isolates belonged to ST25 and ST11 sequence types, while two isolates were assigned to ST105 and ST1212, respectively. We further identified lineages with fastGEAR/BAPS and found that CRKP contained five BAPS lineages with ST11 and ST25 relating to lineages 4 and 5, respectively. The lineage 4 and lineage 5 represented >78% of all isolates and the differences between lineages were consistent with ST types and time collections (Figure.1(a)).

In addition, all ST11 strains harbored *bla_KPC-2_*, whereas *bla_NDM-1_* was found in ST25, ST105 and 3 unknown ST type samples (Figure. 1 (a) (c)). Hierarchical clustering of isolates based on ARG presence or phenotypic susceptibility indicated lineage was the major predictor of resistance-based clustering patterns (Figure.1 (b) (c)).

### Geographical genetic distribution characteristics of *K. pneumoniae*

Through MLST classification of 1579 NGS data from NCBI, a total of 147 known ST types were identified and unknown ST types are found in 384 samples. Among all the known ST type samples, the distribution of epidemic ST varies by counties. Results showed that ST11 (282/1195) and ST258 (264/1195) were dominated, followed by ST15, ST512, ST147 and ST231. In addition, ST11 and ST258 were dominant in China (52%) and the United States (54%), respectively.

Then we detect the presence of *bla_KPC-2_* and *bla_NDM-1_* gene in 1579 NGS genomes and found 462 genomes carried *bla_KPC-2_* gene and 106 had *bla_NDM-1_* gene, respectively. Gene *bla_KPC-2_* was most detected in ST11(88.2%), ST45(80.0%) and ST437 (66.7%) (Figure.3(a)), while *bla_NDM-1_* most found in ST1(66.7%), ST14(53.8%)(Figure.3(a)(c)). Besides, only four genomes contained both genes and they were all isolated from China with three ST11 isolates and one ST86 strain.

Among the dominant ST type, we found 88% of ST11 *K. pneumonia* contained *bla_KPC-2_* and 4% had *bla_NDM-1_*. Whereas in ST258 *K. pneumonia*, only 28% of genomes carried *bla_KPC-2_* gene and none of them contained *bla_NDM-1_*. The presence of *bla_KPC-2_* and *bla_NDM-1_* gene in China (9% and 62%) was significantly higher than those in the United States (2% and 18%) (Figure.2).

**Fig. 2.**
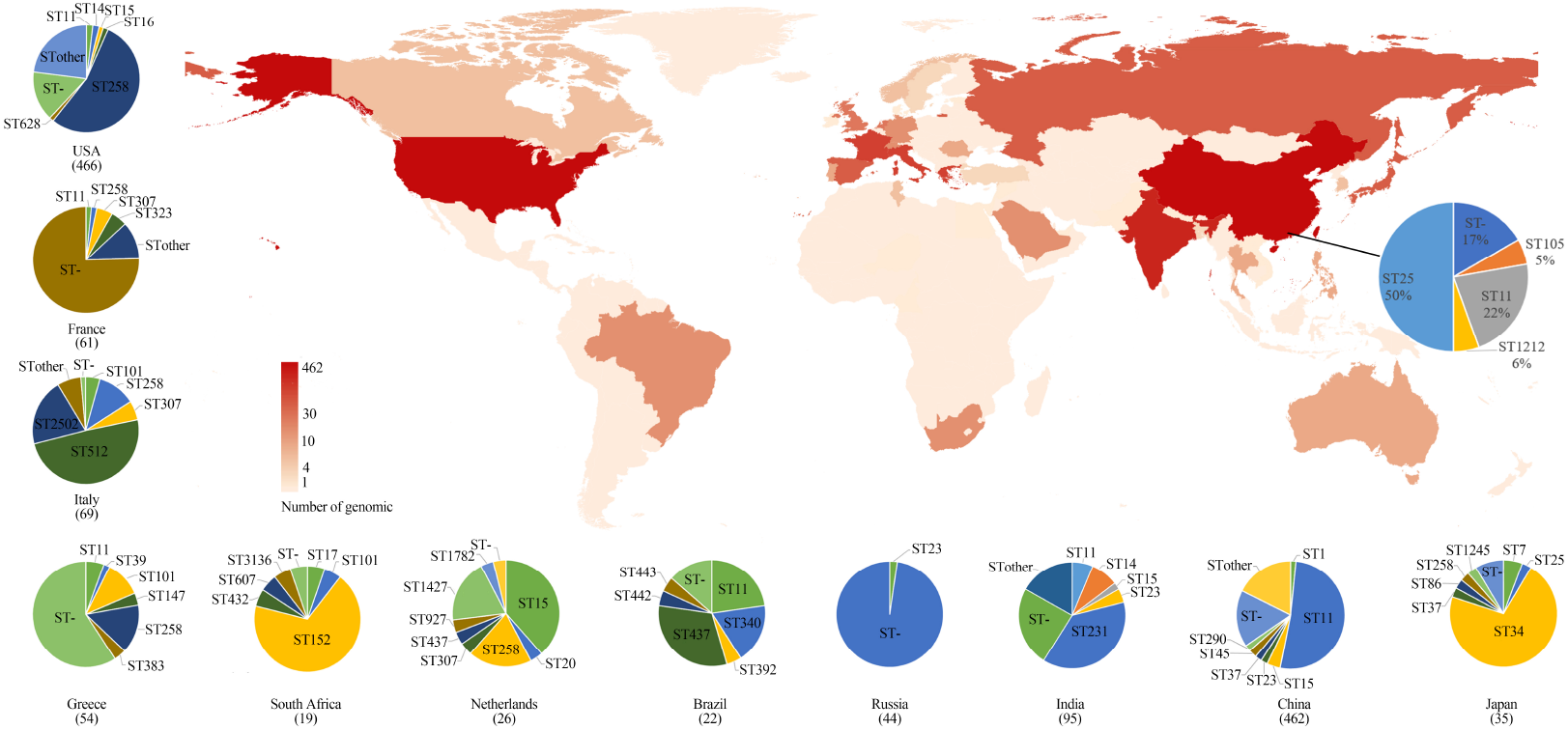
Geographic distribution and MLST typing results of *K. pneumoniae* samples in different countries and regions of genomic samples from NCBI and Shunde hospital; The color shades on the map represent the number of samples. The pie chart link with black line to China on the main map represents the MLST typing results of samples from Shunde hospital. The pie chart next to the main map represent MLST typing results of 13 countries.

**Fig. 3.**
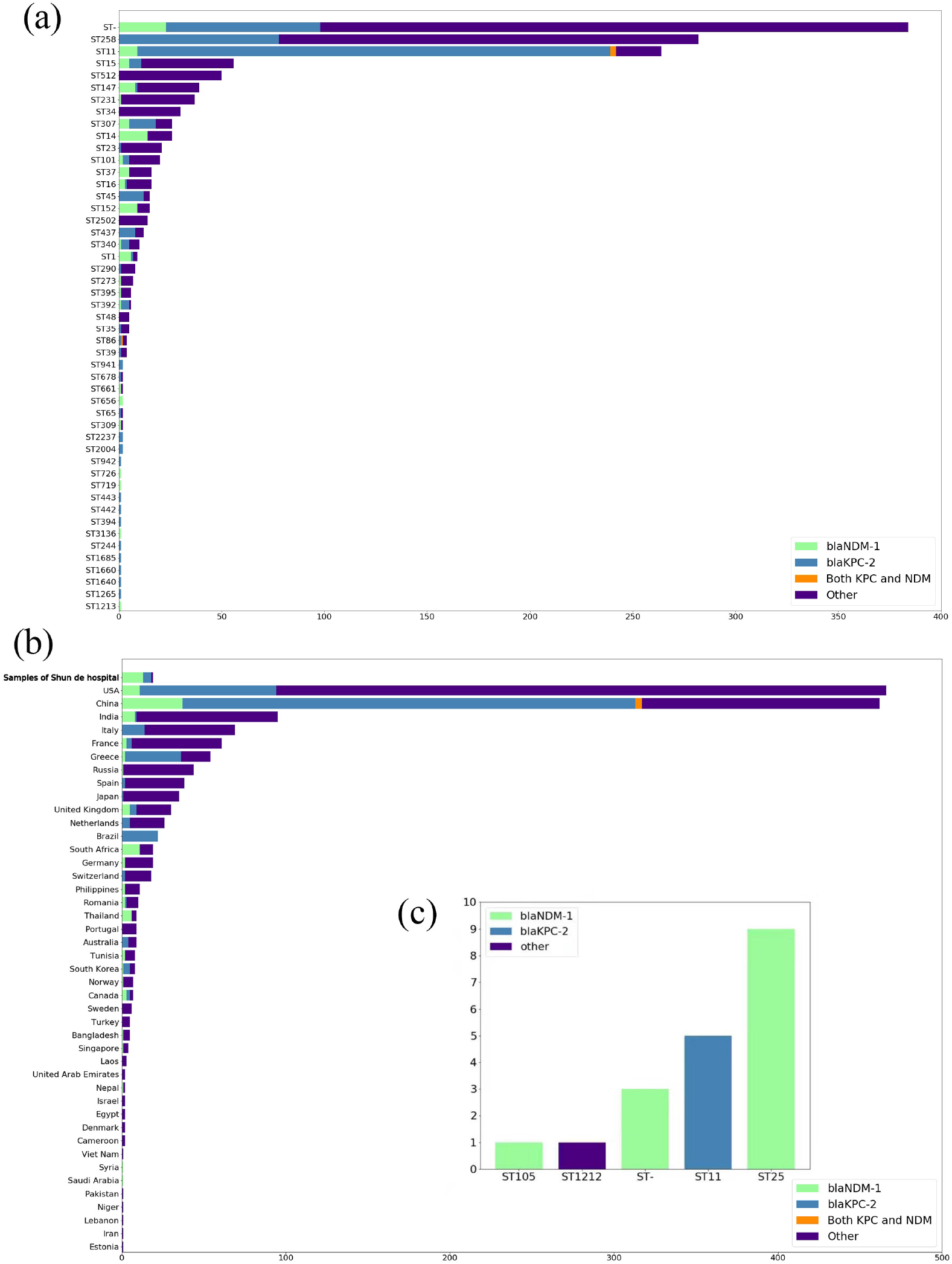
Distribution of *K. pneumoniae* and carbapenem resistance genes (CRGs) (*bla_KPC-2_* and *bla_NDM-1_*); Tpyes of (CRGs) are annotated by different colors. All STs are based on the Pubmlst MLST scheme. (a) Distribution of carbapenem resistance genes of CRGs in *K. pneumoniae* genomes from NCBI in the ST type with large abundance or with *bla_KPC-2_* or *bla_NDM-1_* for display. (b) Geographical distribution of CRGs in *K. pneumoniae* genomes from NCBI (only countries with ≥1 CRG-containing genome are shown). Countries are shown on the y-axis and the numbers on x-axis indicate the number of CRGs. (c) Distribution of carbapenem resistance genes of CRGs in *K. pneumoniae* genomes from Shunde hospital.

### Gene environments of *bla_KPC-2_* and *bla_NDM-1_*

Seventy-six *K. pneumoniae* genomes and thirty *K. pneumoniae* genomes with complete chromosomal genomic data were put into *bla_KPC-2_* and *bla_NDM-1_* gene environment analysis, respectively. For the *bla_KPC-2_* gene, we found 97% (74/76) of *bla_KPC-2_* gene were located on the plasmid. And *bla_KPC-2_* gene is often linked to mobile component *ISKpn6* and *ISKpn27*, where *ISKpn6* is often located in *bla_KPC-2_* gene 3’, and *ISKpn27* often located in *bla_KPC-2_* gene 5’, including those strains belong to ST11 and ST258. However, ISKpn7 was detected in *bla_KPC-2_* gene 5’, which was located in the chromosome of ST258 sample from the United States. However, in ST11 samples from China, the *IS26* family transposon *Tn3* (*tnpR*) is often detected in the gene environment of *bla_KPC-2_*. In addition, aminoglycoside antibiotic resistance gene *accA4* was detected in the *bla_KPC-2_* gene environment of ST258 samples from the United States (Figure.4(a))

**Fig. 4.**
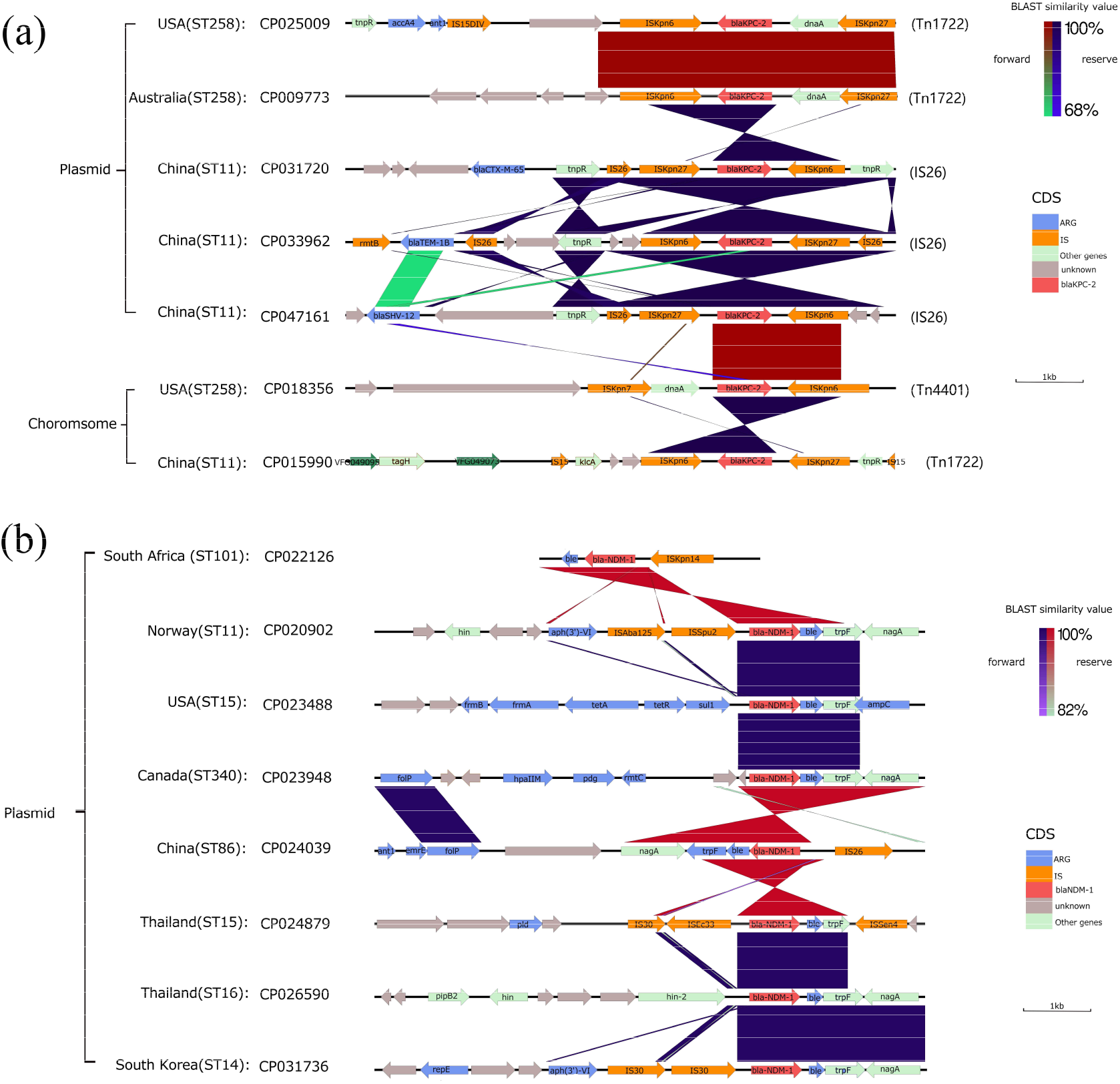
*bla_KPC-2_* and *bla_NDM-1_* gene environment and comparison of some representative samples; BLAST similarity values and the types of different CDS are annotated by different colors next to main chart (a) Gene environment of *bla_KPC-2_* (b) Gene environment of *bla_NDM-1_*

As shown in the Figure.4(b), we found that *bla_NDM-1_* gene was all located on plasmids of all genomes. Gene *bla_NDM-1_* is often linked to *ble*, *trpF* and *nagA* encoding genes for bleomycin resistance protein. In most samples, the *bla_NDM-1_* gene environment contained *IS30* family mobile elements such as ISAba125. Besides, in the ST11 genome from Norway, IS630 family mobile element ISSpu2 and DNA transferase (*hin*) encoding gene fragments were detected in 5’ of *bla_NDM-1_* gene, while in ST16 genome from Thailand, two DNA transferase (*hin*) encoding gene fragments were detected in 5’ of *bla_NDM-1_* gene. In addition, multiple antibiotic resistance genes were detected in the *bla_NDM-1_* gene environment, such as the tetracycline resistance gene (*tet*) and aminoglycoside antibiotic resistance gene (*aph*).

## Discussion

*K. pneumoniae* is responsible for human infections and patient population is the most important reservoir in high-frequency nosocomial *K. pneumoniae* outbreaks. The emergence of *K. pneumonia* carrying carbapenemases and their worldwide dissemination represent significant public health crisis^26^. As carbapenem resistance *bla_KPC-2_* and *bla_NDM-1_* are the most widely reported among CRKP worldwide, we used genomic and phylogenetic approaches to analyze CRKP isolated from hospital in China and uncover the distribution of these two genes in global genomes from GeneBank of NCBI.

Our findings indicate that the CRKP population is highly diverse, encompassing five lineages and various STs. All these CRKP isolates had high AEG burdens and were only sensitive to polymyxins and tigecycline, suggesting resistance in CRKP isolates is more and more serious^27^. In particular, ST11 CRKP have been associated with the global dissemination of *K. pneumoniae*, all the ST types found in hospital were also widely reported in China^28^. Interestingly, through core genome phylogenetic analysis, there was no significant genetic difference in the core genome except the sample P38, which indicated there is high homology between samples in different time of the hospitals. Hence, based on the actual collection time of the samples (at least in different months of the same year), we can infer that there are certain nosocomial infections and clonal transmission in the time segment of the hospital.

MLST typing results showed that ST11-CRKP (5 in 19 strains) in the hospital carried *bla_KPC-2_*. However, according to the genotyping results of GeneBank genome data, 88.25% of ST11 strains contained *bla_KPC-2_* and 4.5% carried *bla_NDM-1_*. Considering the number of samples, these two values are at a relatively high level compared with other ST types. The results of our analysis and previous studies showed that the dominant type of *K. pneumoniae* in China is ST11, which is consistent with previous studies^29,30^. This type of *K. pneumoniae* isolate has obvious preference for carrying *bla_KPC-2_* and *bla_NDM-1_*, which also has geographical differences. Considering the sample size, the proportion of isolates in China carrying the two major carbapenem resistance genes, especially the proportion of *bla_KPC-2_* gene, is at a relatively high level in the world. This also means that isolates prevalent in China are more likely to develop pan-carbapenem drug resistance.

In addition, *bla_KPC-2_* gene was found in ST45, ST307 and ST437 isolates of ST-type *K. pneumoniae*, among which ST437 was the dominant ST-type in Brazil. In general, these three ST types of *K. pneumoniae* were widely distributed in the global scope and had no obvious locality, so it was worth exploring the underlying isolate mobility problem.

Besides, co-occurrence of *bla_NDM-1_ andbla_KPC-2_* in a clinical isolate of *K. pneumoniae* was considered to result in very broad-spectrum antibiotic resistance profiles^31^. Previous studies have found this phenomenon in India, South Africa, and China^32,33^. In our analysis, the strains with co-expression characteristics selected were all from China, which indicated that the detection of multi-drug resistant bacteria in China is relatively adequate, and also indicated that the abuse of antibiotics in China is serious and worthy of vigilance. In addition, all ST25 samples isolated (9 isolates) in our study carried *bla_NDM-1_*. This preference has not been previously reported and is worth further exploration.

It was noted that through genetic analysis we found that the resistant gene *bla_KPC-2_* and adjacent mobile component always form *Tn3-ISKpn27-bla_KPC-2_ -ISKpn6* chain or *ISKpn6- bla_KPC-2_ -ISKpn27-IS26*, which were called *Tn1*722-based unit transposon and IS26-based composite transposon in previous studies, respectively^34–36^. This phenomenon means that the core structure *ISKpn27-bla_KPC-2_ -ISKpn6* of the plasmid is likely to carry *bla_KPC-2_* to transfer in the global scope. In addition, the presence of *bla_KPC-2_* on plasmids was more common in earlier reports, but our study showed that *bla_KPC-2_* in some isolate was located on the chromosome genome, and one of the *bla_KPC-2_* isolates carried *ISKpn7* instead of *ISKpn27* in the core structure. However, the insertion sequence *ISKpn6* and *ISKpn7* were often reported in transposon *Tn4401*, which was the main genetic structure enhancing the spread of the *bla_KPC-_* type genes onto different plasmid scaffolds^37^. In addition, the presence of a variety of carbapenem resistance genes was detected in the *bla_KPC-2_* environment of ST11 samples from China. The interaction between these carbapenem resistance genes and *bla_KPC-2_* and the influence on the drug resistance phenotype are worth further investigation.

Besides, we found that there were *ISAba125*, isomerase *trpF, ISSen4* and family elements in the environment of *bla_NDM-1_* in some isolates, and the upstream and downstream of *bla_NDM-1_* gene often contained transposons (*Tn3*) or inserted sequence fragments (*IS30*), which often involved in horizontal transfer of the drug-resistant gene^38^. In most *bla_NDM-1_* samples, *ble*, *trpF* and *nagA* are closely linked downstream, so it can be considered that *bla_NDM-1_-ble-trpf-nagA* may be the core structure of horizontal transfer of *bla_NDM-1_*^39^.

DNA transferase (*hin*) is a special recombinant binding enzyme that can promote the inversion of DNA position and thus control the alternating expression of proteins and participate in the specific recombination of bacteria, which has been reported to be involved in the inversion control of DNA segment H in the flagellum phase of *Salmonella enterica*^40^. This hint that *hin* may be involved in the horizontal transfer or expression control of *bla_NDM-1_*.

## Conclusion

In this study, we found all CRKP strains had high ARG and antibiotic burdens and belonged to the dominant ST25 and ST11. Phylogenetic analysis showed that there was clonal transmission of CRKP in the hospital. Bioinformatic analysis of 1597 NCBI samples revealed that ST11 and ST258 were the most detected clones and both ST types had a preference for carrying *bla_KPC-2_* gene. The core structure of *ISKpn27-bla_KPC-2_-ISKpn6* and *bla_NDM-1_-ble-trpF-nagA* is highly epidemic in KPC- and NDM-1-producing *K. pneumoniae*. In addition, DNA transferase (*hin*) may play an important role in the horizontal transfer or expression of *bla_NDM-1_* and needed to be further studied. Our study provides a complete genetic background and geographical distribution for further understanding the transmission mode and mechanism of *bla_KPC-2_* and *bla_NDM-1_*.

## Authors’ contributions

The number one author contributed the same as the number two author.

## Acknowledgments

This work was supported by the National Natural Science Foundation of China (Grant No. 31672608) and the National College Students Innovation and Entrepreneurship Foundation of China (201910564054).

## Reference

1. Martin RM, Bachman MA. Colonization, Infection, and the Accessory Genome of *Klebsiella pneumoniae*. Frontiers in cellular and infection microbiology 2018; 8: 4-.doi.

2. Tacconelli E, Carrara E, Savoldi A, et al. Discovery, research, and development of new antibiotics: the WHO priority list of antibiotic-resistant bacteria and tuberculosis. The Lancet Infectious Diseases 2018; 18: 318–27.doi.

3. Sawa T, Kooguchi K, Moriyama K. Molecular diversity of extended-spectrum beta-lactamases and carbapenemases, and antimicrobial resistance. Journal of intensive care 2020; 8: 13.doi.

4. Candan ED, Aksöz N. *Klebsiella pneumoniae*: characteristics of carbapenem resistance and virulence factors. Acta Biochim Pol 2015; 62: 867–74.doi.

5. Munoz-Price LS, Poirel L, Bonomo RA, et al. Clinical epidemiology of the global expansion of *Klebsiella pneumoniae* carbapenemases. The Lancet Infectious diseases 2013; 13: 785–96.doi.

6. Abbas HA, Kadry AA, Shaker GH, Goda RM. Impact of specific inhibitors on metallo-beta-carbapenemases detected in *Escherichia coli* and *Klebsiella pneumoniae* isolates. Microbial pathogenesis 2019; 132: 266–74.doi.

7. Nordmann P, Poirel L. The difficult-to-control spread of carbapenemase producers among *Enterobacteriaceae* worldwide. Clin Microbiol Infect 2014; 20: 821–30.doi.

8. Cerdeira LT, Lam MMC, Wyres KL, et al. Small IncQ1 and Col-Like Plasmids Harboring bla KPC-2 and Non-Tn4401 Elements (NTEKPC-IId) in High-Risk Lineages of *Klebsiella pneumoniae* CG258. Antimicrob Agents Chemother 2019; 63.doi.

9. Abramowicz L, Gerard M, Martiny D, Delforge M, De Wit S, Konopnicki D. Infections due to carbapenemase-producing bacteria, clinical burden, and impact of screening strategies on outcome. Medecine et maladies infectieuses 2020.doi.

10. Ferrari C, Corbella M, Gaiarsa S, et al. Multiple *Klebsiella pneumoniae* KPC Clones Contribute to an Extended Hospital Outbreak. Front Microbiol 2019; 10: 2767.doi.

11. Chen L, Mathema B, Chavda KD, DeLeo FR, Bonomo RA, Kreiswirth BN. Carbapenemase-producing *Klebsiella pneumoniae*: molecular and genetic decoding. Trends in microbiology 2014; 22: 686–96.doi.

12. Pitout JDD, Nordmann P, Poirel L. Carbapenemase-Producing *Klebsiella pneumoniae*, a Key Pathogen Set for Global Nosocomial Dominance. Antimicrobial agents and chemotherapy 2015; 59: 5873–84.doi.

13. Gu D, Dong N, Zheng Z, et al. A fatal outbreak of ST11 carbapenem-resistant hypervirulent *Klebsiella pneumoniae* in a Chinese hospital: a molecular epidemiological study. The Lancet Infectious diseases 2018; 18: 37–46.doi.

14. Du P, Zhang Y, Chen C. Emergence of carbapenem-resistant hypervirulent *Klebsiella pneumoniae*. The Lancet Infectious diseases 2018; 18: 23–4.doi.

15. Fontana C, Angeletti S, Mirandola W, et al. Whole genome sequencing of carbapenem-resistant *Klebsiella pneumoniae*: evolutionary analysis for outbreak investigation. Future Microbiol 2020.doi.

16. Ramsamy Y, Mlisana KP, Allam M, et al. Genomic Analysis of Carbapenemase-Producing Extensively Drug-Resistant *Klebsiella pneumoniae* Isolates Reveals the Horizontal Spread of p18-43_01 Plasmid Encoding bla_NDM-1_ in South Africa. Microorganisms 2020; 8.doi.

17. Clinical and Laboratory Standards Institute. Performance Standards for Antimicrobial Susceptibility Testing, M100. 28th Informational Supplement. Clinical and Laboratory Standards Institute. Wayne, PA; 2018.

18. EUCAST. European Committee on Antimicrobial Susceptibility Testing. Breakpoint Tables for Interpretation of MICs and Zone Diameters. http://www.eucast.org/fileadmin/src/media/PDFs/EUCAST_files/Breakpoint_tables/v_8.0_Breakpoint_Tables.pdf.

19. Goris J, Konstantinidis KT, Klappenbach JA, Coenye T, Vandamme P, Tiedje JM. DNA-DNA hybridization values and their relationship to whole-genome sequence similarities. International journal of systematic and evolutionary microbiology 2007; 57: 81–91.doi.

20. D’Souza AW, Potter RF, Wallace M, et al. Spatiotemporal dynamics of multidrug resistant bacteria on intensive care unit surfaces. Nat Commun 2019; 10: 4569.doi.

21. Maiden MC, Jansen van Rensburg MJ, Bray JE, et al. MLST revisited: the gene-by-gene approach to bacterial genomics. Nat Rev Microbiol 2013; 11: 728–36.doi.

22. Kichenaradja P, Siguier P, Perochon J, Chandler M. ISbrowser: an extension of ISfinder for visualizing insertion sequences in prokaryotic genomes. Nucleic Acids Res 2010; 38: D62–8.doi.

23. Seemann T. Prokka: rapid prokaryotic genome annotation. Bioinformatics (Oxford, England) 2014; 30: 2068–9.doi.

24. Page AJ, Cummins CA, Hunt M, et al. Roary: rapid large-scale prokaryote pan genome analysis. Bioinformatics (Oxford, England) 2015; 31: 3691–3.doi.

25. Mostowy R, Croucher NJ, Andam CP, Corander J, Hanage WP, Marttinen P. Efficient Inference of Recent and Ancestral Recombination within Bacterial Populations. Molecular biology and evolution 2017; 34: 1167–82.doi.

26. Effah CY, Sun T, Liu S, Wu Y. *Klebsiella pneumoniae*: an increasing threat to public health. Ann Clin Microbiol Antimicrob 2020; 19: 1.doi.

27. Gheitani L, Fazeli H, Moghim S, Nasr Isfahani B. Frequency Determination of Carbapenem-Resistant *Klebsiella pneumoniae* (CRKP) Isolated from hospitals in Isfahan of Iran and Evaluation of Synergistic Effect of Colistin and Meropenem on them. Cellular and molecular biology (Noisy-le-Grand, France) 2018; 64: 70–4.doi.

28. Gu B, Bi R, Cao X, Qian H, Hu R, Ma P. Clonal dissemination of KPC-2-producing *Klebsiella pneumoniae* ST11 and ST48 clone among multiple departments in a tertiary teaching hospital in Jiangsu Province, China. Annals of translational medicine 2019; 7: 716.doi.

29. Zhan L, Wang S, Guo Y, et al. Outbreak by Hypermucoviscous *Klebsiella pneumoniae* ST11 Isolates with Carbapenem Resistance in a Tertiary Hospital in China. Frontiers in cellular and infection microbiology 2017; 7: 182.doi.

30. Fu P, Tang Y, Li G, Yu L, Wang Y, Jiang X. Pandemic spread of bla_KPC-2_ among *Klebsiella pneumoniae* ST11 in China is associated with horizontal transfer mediated by IncFII-like plasmids. Int J Antimicrob Agents 2019; 54: 117–24.doi.

31. Kumarasamy K, Kalyanasundaram A. Emergence of *Klebsiella pneumoniae* isolate co-producing NDM-1 with KPC-2 from India. J Antimicrob Chemother 2012; 67: 243–4.doi.

32. Brink AJ, Coetzee J, Clay CG, et al. Emergence of New Delhi metallo-beta-lactamase (NDM-1) and *Klebsiella pneumoniae* carbapenemase (KPC-2) in South Africa. J Clin Microbiol 2012; 50: 525–7.doi.

33. Gao H, Liu Y, Wang R, Wang Q, Jin L, Wang H. The transferability and evolution of NDM-1 and KPC-2 co-producing *Klebsiella pneumoniae* from clinical settings. EBioMedicine 2020; 51: 102599.doi.

34. Wang L, Fang H, Feng J, et al. Complete sequences of KPC-2-encoding plasmid p628-KPC and CTX-M-55-encoding p628-CTXM coexisted in *Klebsiella pneumoniae*. Front Microbiol 2015; 6: 838.doi.

35. Li G, Zhang Y, Bi D, et al. First report of a clinical, multidrug-resistant *Enterobacteriaceae* isolate coharboring fosfomycin resistance gene fosA3 and carbapenemase gene bla_KPC-2_ on the same transposon, Tn1721. Antimicrob Agents Chemother 2015; 59: 338–43.doi.

36. Chen YT, Lin JC, Fung CP, et al. KPC-2-encoding plasmids from Escherichia coli and *Klebsiella pneumoniae in Taiwan*. J Antimicrob Chemother 2014; 69: 628–31.doi.

37. Cuzon G, Naas T, Nordmann P. Functional characterization of Tn4401, a Tn3-based transposon involved in bla_KPC_ gene mobilization. Antimicrobial agents and chemotherapy 2011; 55: 5370–3.doi.

38. Nordmann P, Poirel L, Walsh TR, Livermore DM. The emerging NDM carbapenemases. Trends in microbiology 2011; 19: 588–95.doi.

39. Kutsukake K, Nakashima H, Tominaga A, Abo T. Two DNA invertases contribute to flagellar phase variation in *Salmonella enterica serovar Typhimurium* strain LT2. Journal of bacteriology 2006; 188: 950–7.doi.

40. Feng JA, Johnson RC, Dickerson RE. Hin recombinase bound to DNA: the origin of specificity in major and minor groove interactions. Science (New York, NY) 1994; 263: 348–55.doi.

